# Late Onset Alzheimer’s disease risk variants in cognitive decline: The PATH Through Life Study

**DOI:** 10.1101/067694

**Authors:** Shea J. Andrews, Debjani Das, Kaarin J. Anstey, Simon Easteal

## Abstract

Recent genome wide association studies have identified a number of single nucleotide polymorphisms associated with late onset Alzheimer’s disease. Here we examine the associations of 24 LOAD risk loci, individually and collectively as a genetic risk score, with cognitive function. We used data from 1,626 non-demented older Australians of European ancestry who were examined up to four times over 12 years on tests assessing episodic memory, working memory, vocabulary and information processing speed. Linear mixed models were generated to examine associations between genetic factors and cognitive performance. Twelve SNPs were significantly associated with baseline cognitive performance (*ABCA7, MS4A4E, SORL1*), linear rate of change (*APOE, ABCA7, INPP5D, ZCWPW1, CELF1*) or quadratic rate of change (*APOE, CLU, EPHA1, HLA, INPP5D, FERMT2*). In addition, a weighted GRS was associated with linear rate of change in episodic memory and information processing speed. Our results suggest that a minority of AD related SNPs may be associated with non-clinical cognitive decline. Further research is required to verify these results and to examine the effect of preclinical AD in genetic association studies of cognitive decline. The identification of LOAD risk loci associated with non-clinical cognitive performance may help in screening for individuals at greater risk of cognitive decline.

## Introduction

Late onset Alzheimer’s disease (LOAD), in which patients show clinical symptoms >65 years of age, is the most common form of dementia and the number of individuals with LOAD is expected to triple by 2050 [1]. LOAD has a long preclinical phase that commences decades before the onset of clinical symptoms, which are characterised by progressive degeneration of brain structure and chemistry resulting in gradual cognitive and functional decline [2]. The neuropathological hallmarks of LOAD are aggregation and accumulation of extracellular Amyloid-b peptides into amyloid plaques and accumulation of intraneuronal hyperphosphorylated and misfolded tau into neurofibrillary tangles. Accumulation of amyloid plaques and neurofibrillary tangles prompt the pathogenesis of AD by promoting alterations in lipid metabolism, neuroinflammation, endocytosis and synaptic dysfunction and loss that ultimately leads to neuronal cell death [3,4].

LOAD has a large genetic component, with the heritability estimated to be 60-80% [5]. *Apolipoprotein (APOE) epsilon 4* (*ε4) was the first common genetic variant to be identified [6] and remains the strongest genetic predictor of LOAD. Beyond *APOE*, recent genome-wide association studies (GWAS) and a meta-analysis by the International Genomics of Alzheimer’s Project (IGAP) have identified single nucleotide polymorphisms (SNPs) at 23 loci associated with LOAD (*ABCA7, BIN1, CD2AP, CD33, CLU, CR1, EPHA1, MS4A4A, MS4A4E, MS4A6A, PICALM, HLA-DRB5, PTK2B, SORL1, SLC24A4-RIN3, DSG2, INPP5D, MEF2C, NME8, ZCWPW1, CELF1, FERMT2* and *CASS4*; [7–12]).

The identified LOAD risk loci are clustered in biological pathways that play an important role in disease onset and progression [13] and are involved in the accumulation of the pathological features of LOAD [14]. Furthermore, post-mortem analysis suggests that the neuropathological hallmarks of LOAD progress to varying degrees in individuals without dementia and are associated with cognitive status and nonclinical cognitive decline [15,16]. LOAD risk genes are, therefore, good candidates for investigating potential genetic associations with cognitive performance and decline. How these loci affect normal cognitive function may inform how they influence LOAD onset and progression.

This cross-over effect is exemplified by *APOE*, which is associated with LOAD and has effects on episodic memory, perceptual speed, executive functioning and global cognitive ability [17,18] mediated predominantly by amyloid-β plaques [19]. Association with cognitive decline of the first 11 LOAD risk loci identified by genome-wide association studies (GWAS) are inconsistent [20–27]. Whether the new risk loci identified by IGAP are associated with cognitive decline has yet to be extensively investigated [28–30].

Here, we report associations of the 24 most significant LOAD risk loci with longitudinal change in cognitive performance (based on four neuropsychological outcomes) over 12 years in 1,626 community dwelling older adults. We investigate whether these loci are associated, either individually, or collectively as genetic risk scores (GRS), with: average differences in cognitive performance; rate of cognitive decline; and acceleration of the rate of decline over time.

## Methods

### Participants

Participants of this study are community dwelling older adults who were recruited into the Personality and Total Health (PATH) through life project, a longitudinal study of health and wellbeing. Participants in PATH were sampled randomly from the electoral rolls of Canberra and the neighbouring town of Queanbeyan into one of three cohorts based on age at baseline, the 20+ (20-24), 40+ (40-44) and 60+ (60-64) cohorts. Participants are assessed at 4-year intervals, and data from 4 waves of assessment are available. The background and test procedures for PATH have been described in detail elsewhere [31]. Written informed consent for participation in the PATH project was obtained from all participants according to the ‘National Statement’ guidelines of the National Health and Medical Research Council of Australia and following a protocol approved by the Human Research Ethics Committee of The Australian National University.

In this study, data for the 60+ cohort were used, with interviews conducted in 2001-2002 (n = 2,551), 2005-2006 (n = 2,222), 2009-2010 (n = 1,973), and 2014-2015 (n = 1645), for a total of 12 years of follow-up. Individuals were excluded from analysis based on the following criteria: attendance at only 1 wave (n=309); no available genomic DNA (n=60); *APOE* ε2/ε4 genotype (n=60, to avoid conflation of the ε2 protective and ε4 risk affect); non-European ancestry (n=110); probable dementia at any wave (MMSE < 24; n=82); self-reported medical history of epilepsy, brain tumours or infections, stroke and transient ischemic attacks (n= 450). Missing values in “Education” (total number of years of education, n = 128) were imputed using the ‘missForest’ package in R [32]. This left a final sample of 1,626 individuals at baseline.

### Cognitive Assessment

All participants were assessed at baseline and at each subsequent interview for the following four cognitive abilities: perceptual speed was assessed using the Symbol Digit Modalities Test, which asks the participant to substitute as many digits for symbols as possible in 90 seconds [33]; episodic memory was assessed using the Immediate Recall of the first trial of the California Verbal Learning Test, which involves recalling a list of 16 nouns [34]; working memory was assessed using the Digit Span Backward from the Wechsler Memory Scale, which presents participants with series of digits increasing in length at the rate of one digit per second and asks them to repeat the digits backwards [35]; and vocabulary was assessed with the Spot-the-Word Test, which asks participants to choose the real words from 60 pairs of words and nonsense words [36] Raw cognitive test scores at each wave and Pearson’s correlation between test scores are presented in Supplementary Tables 1 & 2.

### Genotyping

For this study, we used genotype data for 25 SNPs that have been associated with LOAD (Table 1). Genotyping of 11 GWAS LOAD risk SNPs (in the following loci: *ABCA7, BIN1, CD2AP, CD33, CLU, CR1, EPHA1, MS4A4A, MS4A4E, MS4A6A and PICALM*) using TaqMan OpenArray assays has been reported previously [20]. In this study 16 SNPs were selected for genotyping. These included the 12 LOAD GWAS SNPs, which were identified in a meta-analysis of the previous GWAS studies performed by IGAP (in the following loci: *HLA-DRB5, PTK2B, SORL1, SLC24A4-RIN3, DSG2, INPP5D, MEF2C, NME8, ZCWPW1, CELF1, FERMT2* and *CASS4*; [12]). Three were associated with general cognitive function (MIR2113-rs10457441, AKAP6-rs17522122, TOMM40-rs10119; [28]. One was associated as a haplotype with LOAD (FRMD4A-rs2446581; [37]). We used proxy SNPs that were in LD with four (*HLA-DRB5/HLA-DRB1*-rs9271192 [r^2^ = 1], MEF2C-rs190982 [r^2^ = 0.89], *CELF1*-rs10838725 [r^2^ = 0.99] and *CASS4*-rs7274581 [r^2^ = 0.99]) of the SNPs reported by IGAP, as Taqman assays were unavailable [12].

**Table 1:**
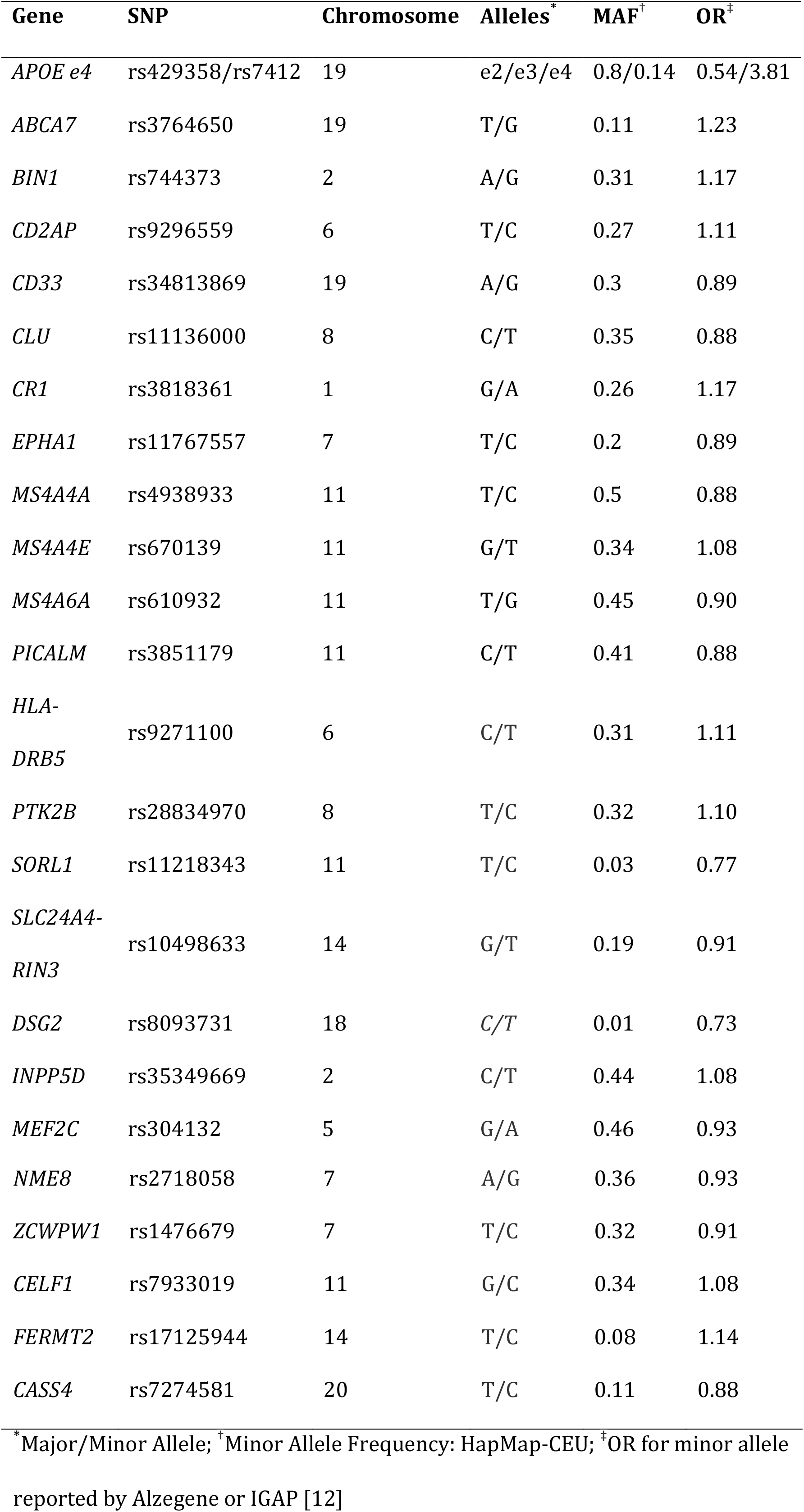
LOAD risk SNPs used in this study

| Gene | SNP | Chromosome | Alleles<sup>*</sup> | MAF<sup>†</sup> | OR<sup>‡</sup> |
| --- | --- | --- | --- | --- | --- |
| *APOE e4* | rs429358/rs7412 | 19 | e2/e3/e4 | 0.8/0.14 | 0.54/3.81 |
| *ABCA7* | rs3764650 | 19 | T/G | 0.11 | 1.23 |
| *BIN1* | rs744373 | 2 | A/G | 0.31 | 1.17 |
| *CD2AP* | rs9296559 | 6 | T/C | 0.27 | 1.11 |
| *CD33* | rs34813869 | 19 | A/G | 0.3 | 0.89 |
| *CLU* | rs11136000 | 8 | C/T | 0.35 | 0.88 |
| *CR1* | rs3818361 | 1 | G/A | 0.26 | 1.17 |
| *EPHA1* | rs11767557 | 7 | T/C | 0.2 | 0.89 |
| *MS4A4A* | rs4938933 | 11 | T/C | 0.5 | 0.88 |
| *MS4A4E* | rs670139 | 11 | G/T | 0.34 | 1.08 |
| *MS4A6A* | rs610932 | 11 | T/G | 0.45 | 0.90 |
| *PICALM* | rs3851179 | 11 | C/T | 0.41 | 0.88 |
| *HLA-DRB5* | rs9271100 | 6 | C/T | 0.31 | 1.11 |
| *PTK2B* | rs28834970 | 8 | T/C | 0.32 | 1.10 |
| *SORL1* | rs11218343 | 11 | T/C | 0.03 | 0.77 |
| *SLC24A4-RIN3* | rs10498633 | 14 | G/T | 0.19 | 0.91 |
| *DSG2* | rs8093731 | 18 | C/T | 0.01 | 0.73 |
| *INPP5D* | rs35349669 | 2 | C/T | 0.44 | 1.08 |
| *MEF2C* | rs304132 | 5 | G/A | 0.46 | 0.93 |
| LOAD genetic risk loci and cognitive decline |  |  |  |  |  |
| *NME8* | rs2718058 | 7 | A/G | 0.36 | 0.93 |
| *ZCWPW1* | rs1476679 | 7 | T/C | 0.32 | 0.91 |
| *CELF1* | rs7933019 | 11 | G/C | 0.34 | 1.08 |
| *FERMT2* | rs17125944 | 14 | T/C | 0.08 | 1.14 |
| *CASS4* | rs7274581 | 20 | T/C | 0.11 | 0.88 |

\*Major/Minor Allele; <sup>†</sup>Minor Allele Frequency: HapMap-CEU; <sup>‡</sup>OR for minor allele reported by Alzegen or IGAP [12]

Genomic DNA was extracted from cheek swabs (n = 2,192) using Qiagen DNA kits or from peripheral blood leukocytes (n = 101) using QIAamp 96 DNA blood kits. Pre-amplification of the targeted loci was performed using the TaqMan PreAmp Master Mix Kit (Life Technologies). Each reaction included 2.5μl TaqMan PreAmp Master Mix (2x), 1.25μl Pre-amplification Assay Pool, 0.5μl H_2_0 and 1.2μl genomic DNA. These reactions were incubated in a Biorad thermocycler for 10 min at 95°C, followed by 12 cycles of 95°C for 15 sec and 60°C for 4 min, and then incubated at 99.9°C for 10 minutes. The PreAmplified products were then held at 4°C until they were diluted 1:20 in 1x TE buffer and then stored at −20°C until use.

Post-PreAmplification, samples were genotyped using the TaqMan OpenArray System. 2μl diluted pre-amplified product was mixed with 2μl TaqMan OpenArray Master Mix. The resulting samples were dispensed using the OpenArray^®^ AccuFill^™^ System onto Format 32 OpenArray plates with each plate containing 96 samples and 16 SNP assays per sample. The QuantStudio^™^ 12K Flex instrument (Applied Biosystems, Carlsbad, California) was used to perform the real-time PCR reactions on the loaded OpenArray plates. The fluorescence emission results were read using the OpenArray^®^ SNP Genotyping Analysis software v1 (Applied Biosystems) and the genotyping analysis was performed using TaqMan^®^ Genotyper v1.3, using the autocalling feature. Manual calls were made on selected genotype calls based on the proximity to the nearest cluster and HapMap positive controls.

Participant-specific quality controls included filters for genotype success rate (> 90%) and sample provenance error assessed via pairwise comparisons of genotype calls between all samples to identify samples with > 90% similarity. Analysis of samples that were flagged in the initial quality control checks were repeated. Those samples that still failed quality control were excluded. SNP-specific filters included genotype call rate (> 90%) and Hardy-Weinberg equilibrium (*p* > 0.05) assessed using an exact test.

The two SNPs defining the *APOE* alleles were genotyped separately using TaqMan assays as previously described [38]. All SNPs were in Hardy-Weinberg equilibrium and genotype frequencies are reported in Supplementary Table 3.

### Data Preparation & Statistical Analysis

All analyses were performed in the R 3.2.3 Statistical computing environment [39]. Cognitive tests at all 4 waves were transformed into z-scores (Mean = 0, SD = 1) using the means and SD at baseline to facilitate comparisons between cognitive tests. A higher score on all tests indicates better cognitive performance.

Genetic dominance was assumed for the previously reported risk allele for all SNPs, except *SORL1, DSG2* and *CASS4* where a recessive model of inheritance was assumed due to the low frequencies of the non-risk allele. The *APOE* *ε2 and *APOE* *ε4 alleles were assumed to be dominant to the *APOE* *ε3 allele. *APOE* alleles were coded as *APOE* *ε2+ (*APOE* *ε2/ε3 + *APOE* *ε2/ε2), APOE*ε4+ (*APOE* *ε4/ε3 + *APOE* *ε4/ε4) or *APOE* *ε3/ε3. Participants with the *APOE* *ε2/ε4 allele were excluded to avoid conflation between the *APOE* *ε2 protective and *APOE* *ε4 risk effects.

Three genetic risk scores were constructed [40]: (1) a simple count genetic risk score (SC-GRS) of the number of risk alleles where 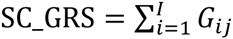; (2) an odds ratio weighted genetic risk score (OR-GRS) where 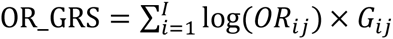; and (3) an explained variance genetic risk score (EV-GRS) weighted by minor allele frequency and odds ratios where 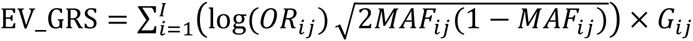. For the above formulae, risk scores are calculated for the ith patient, where log (*OR_i_*) = the odds ratio for the *j*th SNP; *MAF_ij_* = the minor allele frequency for the *j*th SNP; and *G_ij_* = the number of risk alleles for *j*th SNP. SNPs were weighted by their previously reported OR for LOAD and by the minor allele frequency (MAF) reported by the International HapMap project for the CEU reference population (Table 1). Participants missing genetic data for any SNP were excluded from GRS analysis (n = 121). All three GRS were transformed into z-scores to facilitate comparison between them. A higher score indicates greater genetic risk.

Linear mixed effects models (LMMs) with maximum likelihood estimation and subject-specific random intercepts and slopes were used to evaluate the effect of individual SNPs or GRS on longitudinal cognitive performance. Longitudinal change was modelled as a quadratic growth curve, where age centred on baseline was used as an indicator of time; linear rate of change (age) is estimated from the slope of the line tangential to the curve at the intercept and quadratic rate of change (age^2^) is estimated from the acceleration/deceleration in the curve over time. Quadratic growth curves were represented as orthogonal polynomials to avoid collinearity problems and facilitate estimation of the models [41]. Covariates included in the models were sex, total years of education and, for individual SNP models, *APOE* genotype. LMMs were estimated using the R package ‘lme4’ [42]. Statistical significance of the fixed effects was determined using a Kenward-Roger approximation for *F*-tests, where a full model, containing all fixed effects, is compared to a reduced model that excludes an individual fixed effect (R package ‘afex’ [43]). A *p*-value < 0.05 was considered statistically significant. We did not adjust for multiple testing as there is strong a priori evidence for all our hypotheses based on previous findings for validated LOAD susceptibility loci. Conditional 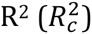, the variance explained by fixed and random effects (i.e. the entire model), and marginal 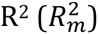, the variance explained by the fixed effects were calculated using the R package ‘MuMIn’ [44–46] by comparing a full model containing the predictor of interest to a reduced model excluding the predictor.

## Results

### Population characteristics of the PATH cohort

Demographic characteristics of the PATH cohort are presented in Table 2. LMMs (Supplementary Table 4) showed that all the cognitive tests were associated with significant linear and quadratic rates of change except for the Digits Span Backwards test. Immediate Recall was associated with linear (β = −22.82; SE = 0.7; *P* = <0.0001) and quadratic rate of change (β = −7.85; SE = 0.69; *P* = <0.0001), with Immediate Recall scores declining with age, and with the decline accelerating over time. Digits Span Backwards Test was associated with linear (β = 1.76; SE = 0.71; *P* = .01) but not quadratic (β = −0.36; SE = 0.66; *P* = .58) rate of change, with Digits Span Backwards test scores increasing with age. Spot-the-Word was associated with linear (β = 4.18; SE = 0.37; *P* = <0.0001) and quadratic (β = −1.84; SE = 0.32; *P* = <0.0001) rate of change, with Spot-the-Word scores increasing with age, and with the rate of change decelerating over time. Symbol Digits Modalities Test was associated with linear (β = −15.36; SE = 0.57; *P* = <0.0001) and quadratic (β = −2.00; SE = 0.51; *P* = <0.0001) rate of change, with Symbol Digits Modalities Test scores declining with age, and with the decline accelerating over time.

**Table 2:** PATH cohort demographics

| Variable | Excluded<br>(n = 924) | Included<br>(n = 1,626) | $t/\chi^2$ | Degrees of<br>Freedom | $P$ |
| --- | --- | --- | --- | --- | --- |
| Age<sup>†</sup> | 62.52 ± 1.51 | 62.49 ± 1.49 | 0.6 | 1938.25 | 0.55 |
| Male<sup>‡</sup> | 831 (51.11) | 485 (52.49) | 0.4 | 1 | 0.53 |
| Education<sup>†</sup> | 14.03 ± 2.57 | 13.31 ± 3.08 | 6.06 | 1649.76 | <0.0001 |
| MMSE<sup>†</sup> | 29.37 ± 0.91 | 28.63 ± 2.08 | 10.23 | 1106.58 | <0.0001 |
| SC-GRS<sup>†</sup> | 24.68 ± 3.21 | 24.6 ± 3.22 | 0.57 | 1322.95 | 0.57 |
| OR-GRS<sup>†</sup> | 3.45 ± 0.81 | 3.46 ± 0.85 | -0.3 | 1256.61 | 0.76 |
| EV-GRS<sup>†</sup> | 1.63 ± 0.41 | 1.64 ± 0.43 | -0.31 | 1260.32 | 0.75 |

<sup>†</sup>unpaired 2-tailed t-test; <sup>‡</sup>Pearson's $\chi^2$ 2-tailed test

Linear rate of change explained 59% - 90% of outcome variation for the entire model, with quadratic rate of change explaining an additional 0.98% - 3.67% of outcome variation. Introducing the covariates into the models explained an additional 8% - 20% of the variation in the fixed effects (Supplementary Table 5).

### Main effects of LOAD GWAS SNPs

Introduction of the 24 LOAD GWAS risk loci into the LMMs (Table 3; for full models including fixed and random effects see Supplementary Tables 6-29) identified 11 SNPs (*APOE, ABCA7, CLU, EPHA1, MS4A4E, HLA, SORL1, INPP5D, ZCWPW1, CELF1, FERMT2*) significantly associated with cognitive performance. The remaining 12 SNPs (*BIN1, CD2AP, CD33, CR1, MS4A4A, MS4A6A, PICALM, PTK2B, SLC24A4-RIN3, DSG2, MEF2C, NME8* and *CASS4*) were not significantly associated with cognitive performance. *APOE* ε4+ was associated with a greater rate of decline in Immediate Recall and Symbol Digit Modalities Tests scores. *APOE* ε4+ and *APOE* ε2+ were both associated with quadratic rate of change in Digits Span Backwards test scores showing a decelerating positive slope. ABCA7-rs3764650-G was associated with a lower initial status at baseline in Immediate Recall Test scores and a reduced rate of decline in Symbol Digit Modalities Test scores. CLU-rs11136000-C was associated with quadratic rate of change in Digits Span Backwards test scores showing an accelerating positive slope. *EPHA1*-rs11767557-T and *HLA*-rs9271100-T were both associated with a reduced rate of growth in Digits Span Backwards test scores. MS4A4E-rs670139-T was associated with increased initial status at baseline in Spot-the-word test scores. *SORL1*-rs11218343-T was associated with a lower initial status at baseline in Symbol Digits Modalities Test scores. INPP5D-rs35349669-T was associated with a reduced rate of decline in Immediate Recall Test scores, a reduced rate of growth in Spot-the-word test scores, a greater rate of decline Symbol Digits Modalities Test scores, and with quadratic rate of change in Digits Span Backwards test scores showing a decelerating positive slope. *ZCWPW1*-rs1476679-T and *CELF1*-rs7933019-C were both associated with an increased rate of growth in Spot-the-word test scores. *FERMT2*-rs17125944-C was associated with a reduced rate of decline in Symbol Digits Modalities Test scores. Comparisons in the R^2^ statistics between covariate only models and the SNPs showed that there was a negligible increase in marginal R^2^ statistics and no increase in conditional R^2^ statistics (Supplementary Tables 6-29).

**Table 3:**
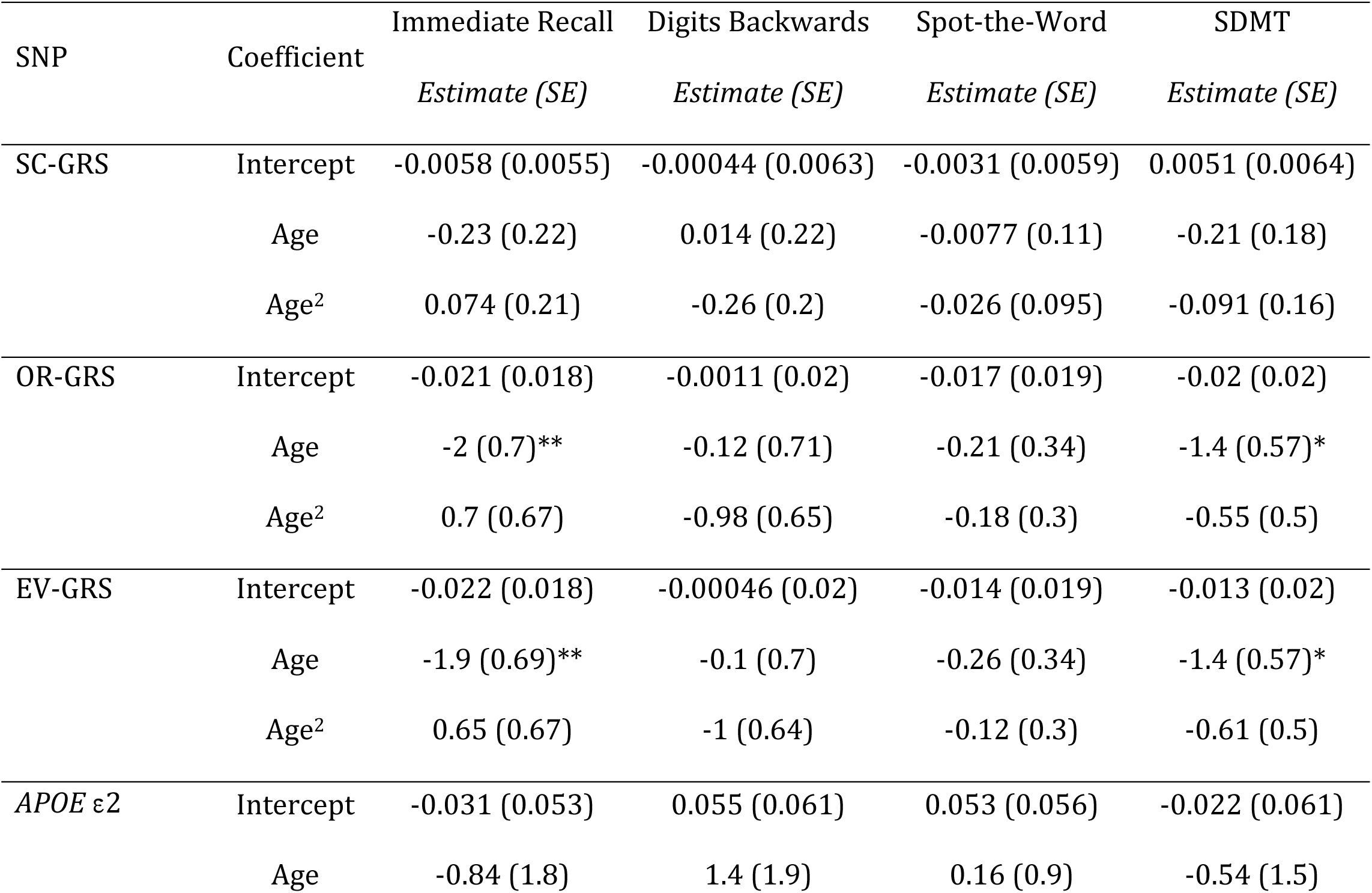
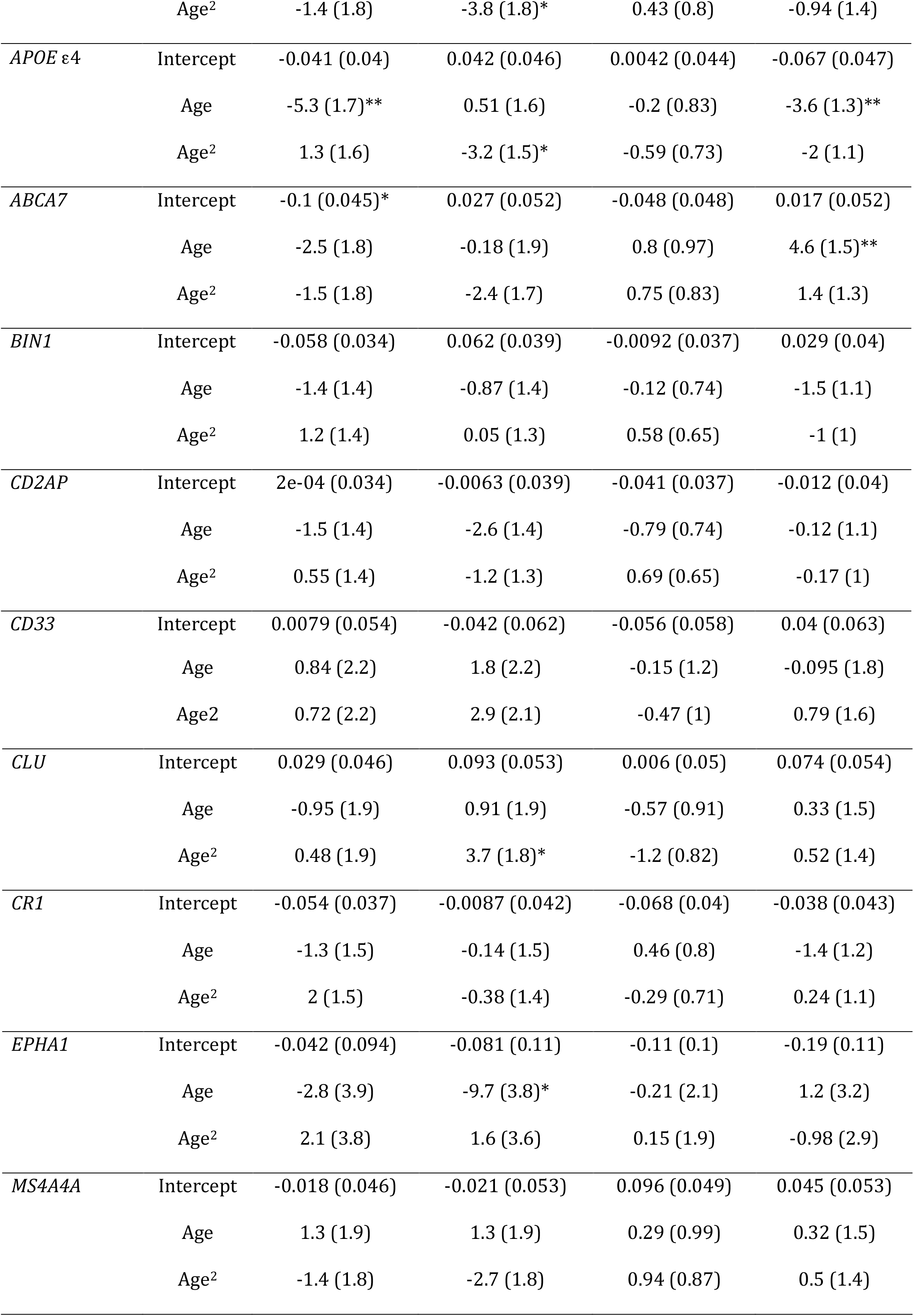
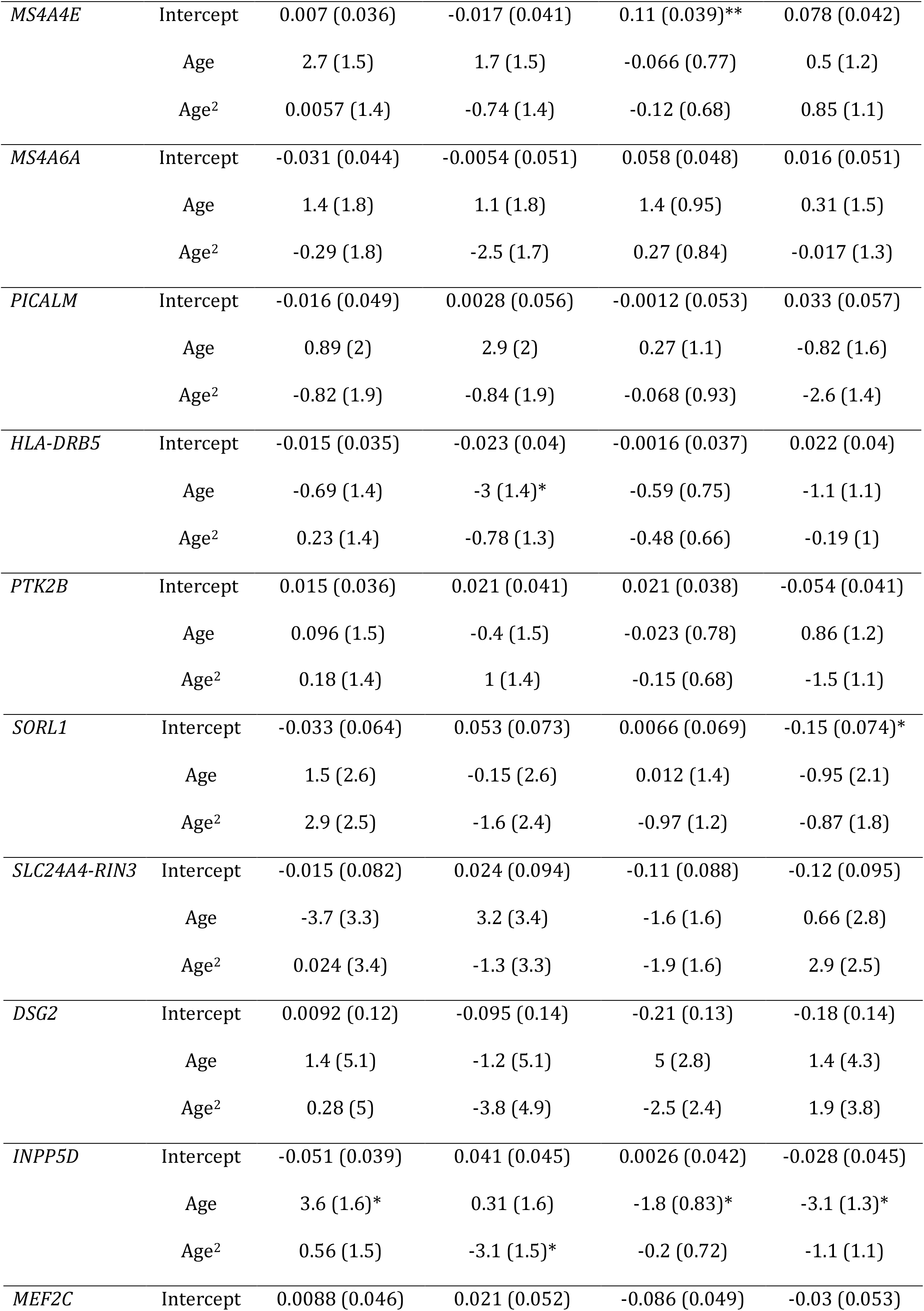
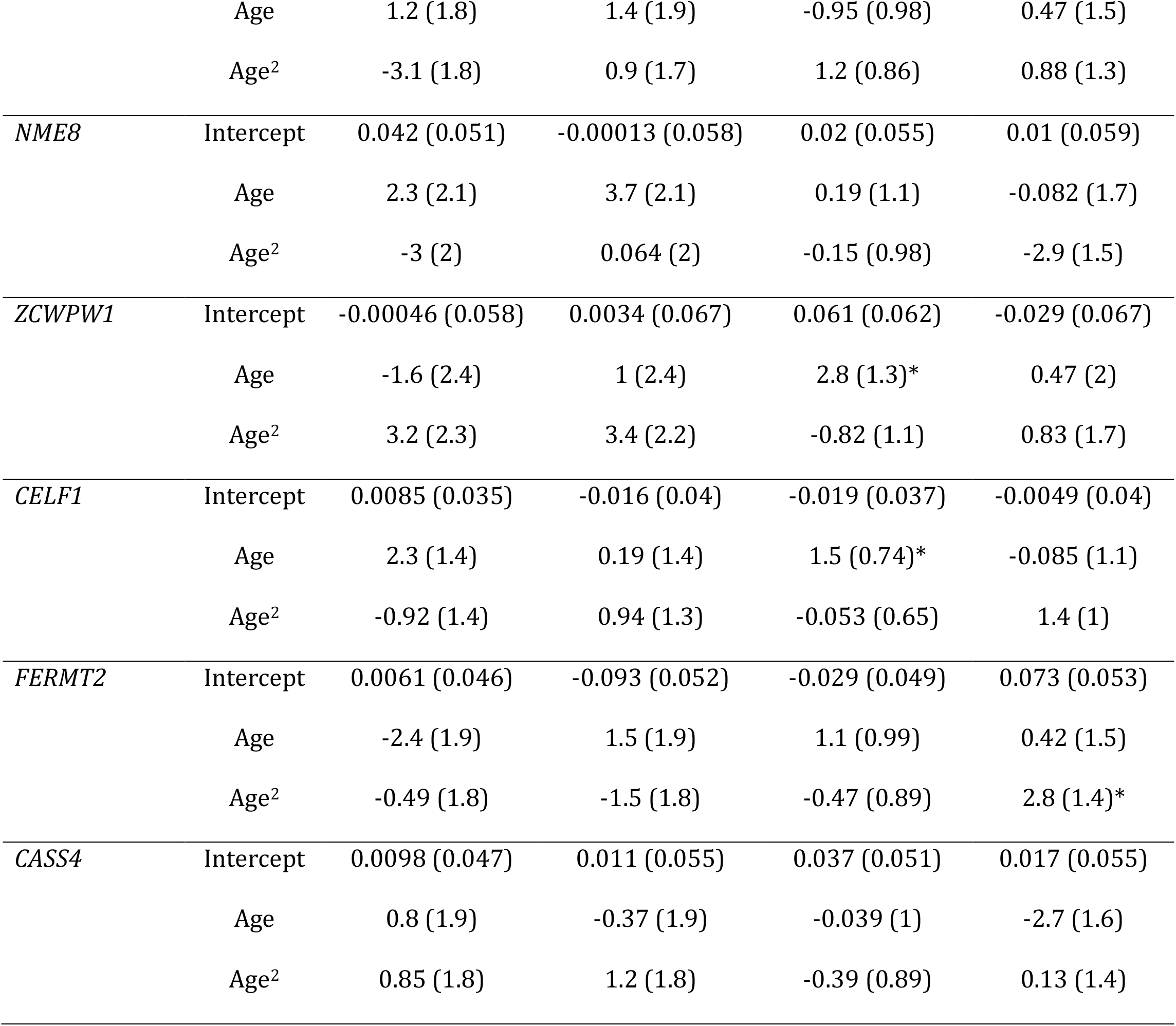
Parameter estimates for the association of LOAD GWAS risk loci with cognitive performance

| SNP | Coefficient | Immediate Recall | Digits Backwards | Spot-the-Word | SDMT |
| --- | --- | --- | --- | --- | --- |
|  |  | *Estimate (SE)* | *Estimate (SE)* | *Estimate (SE)* | *Estimate (SE)* |
| SC-GRS | Intercept | -0.0058 (0.0055) | -0.00044 (0.0063) | -0.0031 (0.0059) | 0.0051 (0.0064) |
|  | Age | -0.23 (0.22) | 0.014 (0.22) | -0.0077 (0.11) | -0.21 (0.18) |
|  | Age<sup>2</sup> | 0.074 (0.21) | -0.26 (0.2) | -0.026 (0.095) | -0.091 (0.16) |
| OR-GRS | Intercept | -0.021 (0.018) | -0.0011 (0.02) | -0.017 (0.019) | -0.02 (0.02) |
|  | Age | -2 (0.7)** | -0.12 (0.71) | -0.21 (0.34) | -1.4 (0.57)* |
|  | Age<sup>2</sup> | 0.7 (0.67) | -0.98 (0.65) | -0.18 (0.3) | -0.55 (0.5) |
| EV-GRS | Intercept | -0.022 (0.018) | -0.00046 (0.02) | -0.014 (0.019) | -0.013 (0.02) |
|  | Age | -1.9 (0.69)** | -0.1 (0.7) | -0.26 (0.34) | -1.4 (0.57)* |
|  | Age<sup>2</sup> | 0.65 (0.67) | -1 (0.64) | -0.12 (0.3) | -0.61 (0.5) |
| *APOE* ε2 | Intercept | -0.031 (0.053) | 0.055 (0.061) | 0.053 (0.056) | -0.022 (0.061) |
|  | Age | -0.84 (1.8) | 1.4 (1.9) | 0.16 (0.9) | -0.54 (1.5) |
|  | Age<sup>2</sup> | -1.4 (1.8) | -3.8 (1.8)* | 0.43 (0.8) | -0.94 (1.4) |
| *APOE ε4* | Intercept | -0.041 (0.04) | 0.042 (0.046) | 0.0042 (0.044) | -0.067 (0.047) |
|  | Age | -5.3 (1.7)** | 0.51 (1.6) | -0.2 (0.83) | -3.6 (1.3)** |
|  | Age<sup>2</sup> | 1.3 (1.6) | -3.2 (1.5)* | -0.59 (0.73) | -2 (1.1) |
| *ABCA7* | Intercept | -0.1 (0.045)* | 0.027 (0.052) | -0.048 (0.048) | 0.017 (0.052) |
|  | Age | -2.5 (1.8) | -0.18 (1.9) | 0.8 (0.97) | 4.6 (1.5)** |
|  | Age<sup>2</sup> | -1.5 (1.8) | -2.4 (1.7) | 0.75 (0.83) | 1.4 (1.3) |
| *BIN1* | Intercept | -0.058 (0.034) | 0.062 (0.039) | -0.0092 (0.037) | 0.029 (0.04) |
|  | Age | -1.4 (1.4) | -0.87 (1.4) | -0.12 (0.74) | -1.5 (1.1) |
|  | Age<sup>2</sup> | 1.2 (1.4) | 0.05 (1.3) | 0.58 (0.65) | -1 (1) |
| *CD2AP* | Intercept | 2e-04 (0.034) | -0.0063 (0.039) | -0.041 (0.037) | -0.012 (0.04) |
|  | Age | -1.5 (1.4) | -2.6 (1.4) | -0.79 (0.74) | -0.12 (1.1) |
|  | Age<sup>2</sup> | 0.55 (1.4) | -1.2 (1.3) | 0.69 (0.65) | -0.17 (1) |
| *CD33* | Intercept | 0.0079 (0.054) | -0.042 (0.062) | -0.056 (0.058) | 0.04 (0.063) |
|  | Age | 0.84 (2.2) | 1.8 (2.2) | -0.15 (1.2) | -0.095 (1.8) |
|  | Age<sup>2</sup> | 0.72 (2.2) | 2.9 (2.1) | -0.47 (1) | 0.79 (1.6) |
| *CLU* | Intercept | 0.029 (0.046) | 0.093 (0.053) | 0.006 (0.05) | 0.074 (0.054) |
|  | Age | -0.95 (1.9) | 0.91 (1.9) | -0.57 (0.91) | 0.33 (1.5) |
|  | Age<sup>2</sup> | 0.48 (1.9) | 3.7 (1.8)* | -1.2 (0.82) | 0.52 (1.4) |
| *CR1* | Intercept | -0.054 (0.037) | -0.0087 (0.042) | -0.068 (0.04) | -0.038 (0.043) |
|  | Age | -1.3 (1.5) | -0.14 (1.5) | 0.46 (0.8) | -1.4 (1.2) |
|  | Age<sup>2</sup> | 2 (1.5) | -0.38 (1.4) | -0.29 (0.71) | 0.24 (1.1) |
| *EPHA1* | Intercept | -0.042 (0.094) | -0.081 (0.11) | -0.11 (0.1) | -0.19 (0.11) |
|  | Age | -2.8 (3.9) | -9.7 (3.8)* | -0.21 (2.1) | 1.2 (3.2) |
|  | Age<sup>2</sup> | 2.1 (3.8) | 1.6 (3.6) | 0.15 (1.9) | -0.98 (2.9) |
| *MS4A4A* | Intercept | -0.018 (0.046) | -0.021 (0.053) | 0.096 (0.049) | 0.045 (0.053) |
|  | Age | 1.3 (1.9) | 1.3 (1.9) | 0.29 (0.99) | 0.32 (1.5) |
|  | Age<sup>2</sup> | -1.4 (1.8) | -2.7 (1.8) | 0.94 (0.87) | 0.5 (1.4) |
| *MS4A4E* | Intercept | 0.007 (0.036) | -0.017 (0.041) | 0.11 (0.039)** | 0.078 (0.042) |
|  | Age | 2.7 (1.5) | 1.7 (1.5) | -0.066 (0.77) | 0.5 (1.2) |
|  | Age<sup>2</sup> | 0.0057 (1.4) | -0.74 (1.4) | -0.12 (0.68) | 0.85 (1.1) |
| *MS4A6A* | Intercept | -0.031 (0.044) | -0.0054 (0.051) | 0.058 (0.048) | 0.016 (0.051) |
|  | Age | 1.4 (1.8) | 1.1 (1.8) | 1.4 (0.95) | 0.31 (1.5) |
|  | Age<sup>2</sup> | -0.29 (1.8) | -2.5 (1.7) | 0.27 (0.84) | -0.017 (1.3) |
| *PICALM* | Intercept | -0.016 (0.049) | 0.0028 (0.056) | -0.0012 (0.053) | 0.033 (0.057) |
|  | Age | 0.89 (2) | 2.9 (2) | 0.27 (1.1) | -0.82 (1.6) |
|  | Age<sup>2</sup> | -0.82 (1.9) | -0.84 (1.9) | -0.068 (0.93) | -2.6 (1.4) |
| *HLA-DRB5* | Intercept | -0.015 (0.035) | -0.023 (0.04) | -0.0016 (0.037) | 0.022 (0.04) |
|  | Age | -0.69 (1.4) | -3 (1.4)* | -0.59 (0.75) | -1.1 (1.1) |
|  | Age<sup>2</sup> | 0.23 (1.4) | -0.78 (1.3) | -0.48 (0.66) | -0.19 (1) |
| *PTK2B* | Intercept | 0.015 (0.036) | 0.021 (0.041) | 0.021 (0.038) | -0.054 (0.041) |
|  | Age | 0.096 (1.5) | -0.4 (1.5) | -0.023 (0.78) | 0.86 (1.2) |
|  | Age<sup>2</sup> | 0.18 (1.4) | 1 (1.4) | -0.15 (0.68) | -1.5 (1.1) |
| *SORL1* | Intercept | -0.033 (0.064) | 0.053 (0.073) | 0.0066 (0.069) | -0.15 (0.074)* |
|  | Age | 1.5 (2.6) | -0.15 (2.6) | 0.012 (1.4) | -0.95 (2.1) |
|  | Age<sup>2</sup> | 2.9 (2.5) | -1.6 (2.4) | -0.97 (1.2) | -0.87 (1.8) |
| *SLC24A4-RIN3* | Intercept | -0.015 (0.082) | 0.024 (0.094) | -0.11 (0.088) | -0.12 (0.095) |
|  | Age | -3.7 (3.3) | 3.2 (3.4) | -1.6 (1.6) | 0.66 (2.8) |
|  | Age<sup>2</sup> | 0.024 (3.4) | -1.3 (3.3) | -1.9 (1.6) | 2.9 (2.5) |
| *DSG2* | Intercept | 0.0092 (0.12) | -0.095 (0.14) | -0.21 (0.13) | -0.18 (0.14) |
|  | Age | 1.4 (5.1) | -1.2 (5.1) | 5 (2.8) | 1.4 (4.3) |
|  | Age<sup>2</sup> | 0.28 (5) | -3.8 (4.9) | -2.5 (2.4) | 1.9 (3.8) |
| *INPP5D* | Intercept | -0.051 (0.039) | 0.041 (0.045) | 0.0026 (0.042) | -0.028 (0.045) |
|  | Age | 3.6 (1.6)* | 0.31 (1.6) | -1.8 (0.83)* | -3.1 (1.3)* |
|  | Age<sup>2</sup> | 0.56 (1.5) | -3.1 (1.5)* | -0.2 (0.72) | -1.1 (1.1) |
| *MEF2C* | Intercept | 0.0088 (0.046) | 0.021 (0.052) | -0.086 (0.049) | -0.03 (0.053) |
|  | Age | 1.2 (1.8) | 1.4 (1.9) | -0.95 (0.98) | 0.47 (1.5) |
|  | Age<sup>2</sup> | -3.1 (1.8) | 0.9 (1.7) | 1.2 (0.86) | 0.88 (1.3) |
| *NME8* | Intercept | 0.042 (0.051) | -0.00013 (0.058) | 0.02 (0.055) | 0.01 (0.059) |
|  | Age | 2.3 (2.1) | 3.7 (2.1) | 0.19 (1.1) | -0.082 (1.7) |
|  | Age<sup>2</sup> | -3 (2) | 0.064 (2) | -0.15 (0.98) | -2.9 (1.5) |
| *ZCWPW1* | Intercept | -0.00046 (0.058) | 0.0034 (0.067) | 0.061 (0.062) | -0.029 (0.067) |
|  | Age | -1.6 (2.4) | 1 (2.4) | 2.8 (1.3)* | 0.47 (2) |
|  | Age<sup>2</sup> | 3.2 (2.3) | 3.4 (2.2) | -0.82 (1.1) | 0.83 (1.7) |
| *CELF1* | Intercept | 0.0085 (0.035) | -0.016 (0.04) | -0.019 (0.037) | -0.0049 (0.04) |
|  | Age | 2.3 (1.4) | 0.19 (1.4) | 1.5 (0.74)* | -0.085 (1.1) |
|  | Age<sup>2</sup> | -0.92 (1.4) | 0.94 (1.3) | -0.053 (0.65) | 1.4 (1) |
| *FERMT2* | Intercept | 0.0061 (0.046) | -0.093 (0.052) | -0.029 (0.049) | 0.073 (0.053) |
|  | Age | -2.4 (1.9) | 1.5 (1.9) | 1.1 (0.99) | 0.42 (1.5) |
|  | Age<sup>2</sup> | -0.49 (1.8) | -1.5 (1.8) | -0.47 (0.89) | 2.8 (1.4)* |
| LOAD genetic risk loci and cognitive decline |  |  |  |  |  |
| *CASS4* | Intercept | 0.0098 (0.047) | 0.011 (0.055) | 0.037 (0.051) | 0.017 (0.055) |
|  | Age | 0.8 (1.9) | -0.37 (1.9) | -0.039 (1) | -2.7 (1.6) |
|  | Age<sup>2</sup> | 0.85 (1.8) | 1.2 (1.8) | -0.39 (0.89) | 0.13 (1.4) |

### Main effects of LOAD GRS

We evaluated the association of three weighted genetic risk scores with cognitive performance (Table 3; Supplementary Tables 30-32). Mean and SD for the raw GRS at baseline are presented in Table 1. The SC-GRS was not associated with cognitive performance. Higher OR- and EV-GRS were associated with a greater rate of decline in Immediate Recall and Symbol Digit Modalities Test scores.

Comparisons in the R^2^ statistics between covariates-only models and the GRS models showed that there was a negligible increase in marginal R^2^ statistics and no increase in conditional R^2^ statistics (Supplementary Tables 30-32). OR- and EV-GRS were not associated with cognitive performance when *APOE* was excluded from the GRS (Supplementary Tables 33-35).

## Discussion

In this study we investigated the association of the 24 most significant LOAD GWAS risk loci with cognitive performance in episodic memory, vocabulary, working memory and processing speed. We identified 11 SNPs as associated with baseline cognitive performance (*ABCA7, MS4A4E, SORL1*), linear rate of change (*APOE, ABCA7, EPHA1, HLA, INPP5D, ZCWPW1, CELF1*) or quadratic rate of change (*APOE, CLU, INPP5D, FERMT2*). GRS, weighted by odds ratio and by odds ratio plus minor allele frequency were both associated with a linear rate of change in episodic memory and processing speed. When *APOE* was excluded from these scores neither GRS were significantly associated with cognitive performance indicating that the association was driven by the dominant effect of the *APOE* *ε4 allele. It should be noted, however, that the effect sizes for the observed associations are small, with an increase in marginal R^2^ statistics ranging from 0.1–0.4% after inclusion of the genetic predictors. In comparison, inclusion of the covariate education in the model increases the marginal R^2^ statistic around 4.3–19.8%.

Previous studies of association between the initial GWAS LOAD risk loci and cognitive performance are characterized by a lack of consistent findings. The limited number of studies that have examined the role of the IGAP LOAD risk loci in cognitive performance also produced mixed results [21–30,47–55].

In univariate analysis, SNPs from 7 of the 23 *non-APOE* GWAS loci have been associated with cognitive performance. *ABCA7* has been associated with declines in the MMSE score in women [30]. *BIN1* has been associated with decline in MMSE score in one study [26]. *CD2AP* has been associated with a composite episodic memory in one study [48]. *CD33* has been associated with a composite executive function score [48] and decline in MMSE in women [30]. *CLU* has been associated with cognitive performance in four studies, with baseline episodic memory [21], baseline and decline in a composite cognitive score [27,53] and decline in 3MS [56]. *CR1* has been associated with declines in verbal fluency [26], global cognition [25,52], episodic memory, perceptual speed, semantic memory [54] and attention [56]. *PICALM* has been associated with a composite cognitive score [53] and decline in global cognition [24]. *NME8* was associated with declines in Clinical Dementia Rating Scale Sum of Boxes Scores [55].

In contrast to a univariate approach, aggregating SNP variation across genomic regions in a ‘gene based’ approach, has identified further AD risk loci as associated with cognitive performance. In a meta-analysis of 31 studies (n = 53,949), *PICALM, MEF2C* and *SLC24A4-RIN3* gene regions were associated with general cognitive function (p ≤0.05). In single sex cohorts, *BIN1, CD33, CELF1, CR1, HLA* cluster, and *MEF2C* gene regions were associated with decline in *MMSE* in a all female cohort and *ABCA7, HLA* cluster, *MS4A6E, PICALM, PTK2B, SLC24A4*, and *SORL1* gene regions were associated with decline in 3MS in a all-male cohort.

Genetic risk scores can have greater predictive power than individual variants because they are based on the cumulative effect of many variants that individually may have effects that are too small to be reliably detected in a univariate analysis. GRS composed of genome wide significant LOAD SNPs identified in the initial LOAD GWAS have been associated with baseline general cognition [48], episodic memory [21], visual memory and MMSE [26] and with decline in episodic memory [21], verbal fluency, visual memory and MMSE [26]. However these associations were no longer statistically significant when *APOE* was excluded from the GRS.

Two studies have investigated a GRS composed of the IGAP LOAD SNPs, one of which showed that a GRS with *APOE* excluded was associated with decline in MMSE in participants with MCI [29]. The second study showed that a GRS with *APOE* included was associated with memory performance at baseline and with a faster rate of decline that accelerated with age. However, after excluding *APOE* only linear rate of change remained significant [57].

Genome-wide significant IGAP LOAD risk loci do not reflect the full spectrum of genetic susceptibility to LOAD risk loci, explaining only 30.62% of the genetic variance of LOAD [58]. Thus, an alternative approach is to construct a genome-wide polygenic score (GPS) which is calculated based not solely on genome-wide significant SNPs, but on all nominally associated variants at a given significance level. The first study to use this method did not find an association with cognitive ability or cognitive change [59]. A more recent study using data collected from the UK Biobank (n = 112 151) found that an AD GRS constructed from 20,437 SNPs that were associated with AD at a threshold of p < 0.05 in the IGAP study was significantly associated with lower verbal-numerical reasoning, memory and educational attainment [60].

As noted above, the association of LOAD risk loci with cognitive performance is mixed. Several factors may explain the discrepancies between studies. First, the failure to replicate positive results between studies could result from differences in participant characteristics (e.g., baseline education, mean age, gender, and ethnicity) and methodologies (e.g., sample size, duration of the study, number of follow-ups, nonlinear time, population stratification, variation in classification, and cognitive measures [61]. In particular, studies that did not exclude cognitively impaired individuals from the analysis could bias the observed results in favour of a positive association [28,62].

Selectively removing from the analysis individuals who develop cognitive impairment during the study, as in this study, may not resolve the issue because of inadvertent inclusion of participants with preclinical dementia. This issue was highlighted in the study by Hassenstab *et al*, which reported that inclusion of individuals who were cognitively normal, but had biomarker and neuroimaging evidence of preclinical AD, greatly exaggerated age-related cognitive decline across multiple cognitive domains [63]. This finding suggests that AD-related genes may be associated with cognitive decline in participants who are in the preclinical stages of AD, and who if followed for long enough would develop AD. This has been observed in cognitively normal participants who had low levels of PET Aβ but were *APOE* *ε4+. This group remained cognitively stable, in comparison to an *APOE* *ε4+ group with high PET Aβ who experienced faster rates of cognitive, decline, suggesting that declines in cognitive function observed in *APOE* *ε4 carriers reflects the effect of *APOE* exacerbating Ab related cognitive decline rather then an independent *APOE* effect[64]. This effect is further indicated by previous studies showing that *ABCA7, EPAH1* and *CLU* were associated with cognitive decline in participants classified as cognitively impaired or demented, but not in those who remained cognitively normal [20,21,51].

Second, the rationale for including LOAD risk loci in the analysis is that they may be associated with biological processes, such as neurotic plaque or neurofibrillary tangle burden, that affect both LOAD and general cognitive performance. Only some LOAD risk loci are known to be associated with these pathological features. There may be a lack of association with cognitive performance because some loci are associated with LOAD for other reasons that do not affect general cognitive function.

Notably, of the 23 loci identified in the IGAP study, only 11 have been associated with neurotic plaque (*ABCA7, BIN1, CASS4, MEF2C, PICALM, MS4A6A, CD33* and *CR1*) or neurofibrillary tangle (*ABCA7, BIN1, CASS4, MEF2C, PICALM, CLU, SORL1* and *ZCWPW1*) burdens in AD case/control autopsies [52,65]. In a longitudinal study, only *BIN1* and *CASS4* were associated with amyloid accumulation [66]. In contrast, in subjects with MCI, none of the LOAD risk loci were associated with levels of Aβ in the cerebrospinal fluid (CSF) and only *SORL1* was associated with levels of tau and phosphorylated tau (components of neurofibrillary tangles) in the CSF.

Furthermore, neurotic plaques and neurofibrillary tangles only explain 30% of the variation in cognitive decline, with cerebrovascular and Lewy body disease neuropatholgies explaining an additional 10% of variation [67]. Given that LOAD neuropathology explains only a small portion of the variation in cognitive decline, the effect sizes of individual LOAD risk loci that influence cognitive decline via amyloid and tau pathways, are expected to be small. This highlights that while LOAD pathology is an important factor in cognitive decline, it occurs in conjunction with other pathological features.

Genetic and environmental factors do not act independently of each other but are likely to interact with each other such that environmental exposures may have differential effects that depend on individual genetic risks and vice versa. Investigating interactions between genetic and environmental and lifestyle risk factors may provide promising results. Interactions between the non-*APOE* risk loci and environmental factors have yet to be extensively investigated, although associations have been observed between *AD* risk loci and physical activity, diabetes, and Mediterranean diet [68–71].

Finally, the pathogenesis of LOAD spans decades, clinically progressing through the preclinical, MCI and dementia stages. The underlying pathological process has been modelled as a cascade that starts with amyloidosis followed by hyperphosphorylated tau accumulation and subsequent structural, functional and cognitive declines [2]. As such, where and when a risk locus is involved in the LOAD pathogenesis cascade may influence whether it is associated with processes that predispose, initiate or propagate cognitive decline. Therefore, the lack of associations with cognitive performance with the majority of the LOAD risk loci in this study could indicate that they exert their pathological effects at latter stages. Associations have been reported between LOAD risk loci *CD2AP, CLU, MS4A6A* and *INPP5D* and progression from normal cognition to dementia [72]; between *CLU, CR1*, and *NME8* and progression from MCI to dementia [72–74]; between *INPP5D, MEFC2, EPHA1, PT2KB, FERMT2, CASS4* and rate of progression in AD [75]; and between *PICALM* and *MS4A6A* and progression to MCI/Dementia from normal cognition normal [21].

In the context of atypical AD, which is characterized by the development of non-amnestic cognitive deficits [76], some LOAD SNPs may be associated with non-amnestic cognitive domains. As such, the lack of consensus across studies could result from participants being at different stages in the pathogenesis of AD and from assessment of different cognitive domains between studies.

The present findings need to be interpreted with an understanding of their limitations. First, the PATH cohort is better educated then the population it was drawn from. As higher education is associated with a reduced risk of cognitive decline and incident dementia, this may limit our ability to detect an association between genetic factors and cognitive performance. Second, the subjects in this study were of European ancestry, and thus the results presented may not be generalizable to other populations. Third, even after retrospectively removing participants with cognitive impairment, this cohort may still contain individuals who are in the preclinical stages of AD and thus potentially biasing our results. Fourth, there may have been differential attrition from the PATH study of individuals who later became severely impaired and demented, which may have biased results because these individuals would not be excluded from our analysis and are more likely to experience faster rates of cognitive decline [77]. Finally, we had a prior expectation that each of the SNPs we analysed would be associated with cognitive decline because of its association with LOAD. We, therefore, did not correct for the effect of multiple comparisons. If we had applied a Bonferroni correction none of the observed associations would remain significant.

Despite these limitations, this study has a number of strengths. It was performed in a large community-based cohort that has been followed for a period of 12 years with four waves of data assessing four separate cognitive domains. This allows for robust statistical modelling of the association of genetic factors with non-linear declines across a broad spectrum of cognition functions. Additionally, the narrow age range of this cohort reduces the influence of age differences on the results.

In conclusion, our results suggest that a subset of AD-risk loci are associated with non-clinical cognitive decline, although the effect size of each locus is small. Further, when demographic and lifestyle factors are taken into account, neither individual SNPs nor GRS explain a significant proportion of the variance in cognitive decline in our sample. Further investigation of the association of LOAD risk loci with cognitive function needs to account for the inclusion of participants with preclinical AD. The use of neuroimaging and cerebrospinal fluid biomarkers to determine preclinical AD status will allow for a more robust analysis of the role of LOAD risk loci in cognitive aging.

## Acknowledgments

We thank the participants of the PATH study, Peter Butterworth, Andrew Mackinnon, Anthony Jorm, Bryan Rodgers, Helen Christensen, Patricia Jacomb and Karen Mawell. The study was supported by the National Health and Medical Research Council (NHMRC) grants 973302, 179805, and 1002160 and the NHMRC Dementia Collaborative Research Centres. DD is funded by NHMRC Project Grant No. 1043256. KJA is funded by NHMRC Research Fellowship No. 1002560.

